# Intact Drosophila Central Nervous System Cellular Quantitation reveals Sexual Dimorphism

**DOI:** 10.1101/2021.11.03.467146

**Authors:** Wei Jiao, Gard Spreemann, Evelyne Ruchti, Soumya Banerjee, Samuel Vernon, Ying Shi, R. Steven Stowers, Kathryn Hess, Brian D. McCabe

## Abstract

Establishing with precision the quantity and identity of the cell types of the brain is a prerequisite for a detailed compendium of gene and protein expression in the central nervous system. Currently however, strict quantitation of cell numbers has been achieved only for the nervous system of *C.elegans*. Here we describe the development of a synergistic pipeline of molecular genetic, imaging, and computational technologies designed to allow high-throughput, precise quantitation with cellular resolution of reporters of gene expression in intact whole tissues with complex cellular constitutions such as the brain. We have deployed the approach to determine with exactitude the number of functional neurons and glia in the entire intact larval *Drosophila* central nervous system (CNS), revealing fewer neurons and more glial cells than previously predicted. We also discover an unexpected divergence between the sexes at this juvenile developmental stage, with the female CNS having significantly more neurons than that of males. Topological analysis of our data establishes that this sexual dimorphism extends to deeper features of CNS organisation. We additionally extended our analysis to quantitate the expression of voltage-gated potassium channel family genes throughout the CNS and uncover substantial differences in abundance. Our methodology enables robust and accurate quantification of the number and positioning of cells within intact organs, facilitating sophisticated analysis of cellular identity, diversity, and gene expression characteristics.

## INTRODUCTION

Establishing the precise number of cells in the brain is essential to create organ-wide catalogues of cell-types and their gene expression (Lent et al., 2012; Devor et al., 2013). However, apart from the nervous system of the nematode *Caenorhabditis elegans* [302 neurons, 56 glia] (White et al., 1986), the exact numbers of cells within the central nervous system (CNS) of model organisms or that of humans is currently unknown, with estimates, including those based upon extrapolation from direct quantification of brain sub-regions, varying widely (Silbereis et al., 2016; Keller et al., 2018; von Bartheld et al., 2016).

Studies of the CNS of *Drosophila melanogaster*, which in scale and behavioural repertoire has been viewed as intermediate between nematodes and rodents (Bellen et al., 2010; Alivisatos et al., 2012), currently include large-scale efforts to establish both a neuronal connectome and cell atlas (Scheffer and Meinertzhagen, 2019; Allen et al., 2020; Li et al., 2022). Nonetheless, the precise numbers of cells (neurons or glia) in either the smaller larval or larger adult *Drosophila* CNS, comprised of both a brain and Ventral Nerve Cord (VNC), remain unknown, though several approximations have been suggested. For the larval CNS, a range of between 10,000 to 15,000 active neurons has been proposed (Scott et al., 2001; Meinertzhagen, 2018; Eschbach and Zlatic, 2020) across developmental timepoints. For adult Drosophila, approximations have been suggested in the range of between 100,000 to 199,000 neurons in the brain (Simpson, 2009; Chiang et al., 2011; Kaiser, 2015; Scheffer and Meinertzhagen, 2019; Raji and Potter, 2021) together with between 10,000 to 20,000 cells in the VNC (Birkholz et al., 2015; Lacin et al., 2019; Bates et al., 2019; Allen et al., 2020). The other major CNS cell type, glia, has been estimated to be approximately 10% of the number of neurons (Kremer et al., 2017; Meinertzhagen, 2018; Raji and Potter, 2021). Given the large diversity of these estimates, precise quantification of the numbers of *Drosophila* neurons and glia would seem a desirable goal, beginning with the smaller larval CNS which enables the wide compendium of larval *Drosophila* behaviours (Gerber et al., 2009; Neckameyer and Bhatt, 2016; Eschbach and Zlatic, 2020; Louis, 2020; Gowda et al., 2021).

Complicating the aspiration quantitate the *Drosophila* larval CNS, in addition to the general problem of separating and quantifying primary cell types such as neurons and glia, are two specific confounding factors that limit simple total cell quantification approaches. Firstly, encompassed within and surrounding the larval CNS are dividing neuroblasts, which will give rise to adult neurons (Doe, 2017). Relatedly, imbedded within the larval CNS are substantial numbers of immature adult neurons, observed from electron micrograph reconstructions as having few or no dendrites and axons that terminate in filopodia lacking synapses (Eichler et al., 2017). These immature neurons are unlikely to contribute to larval CNS function and are generally excluded when considering larval neuronal circuit architecture (Eichler et al., 2017; Scheffer and Meinertzhagen, 2019). It has been suggested that only a small fraction of the total number of larval CNS cells may actually contribute to CNS function (Ravenscroft et al., 2020).

Here we have sought to develop a synergistic molecular genetic, imaging, and computational pipeline designed *de novo* to allow automated neuron, glia, or other gene expression features to be precisely quantitated with cellular resolution in intact whole central nervous systems. Central to the approach are high signal-to-noise gene expression reporters that produce a punctate, nucleus-localised output facilitating downstream automated computational measurement and analysis. Exploiting multiple genetic reagents designed to selectively identify only functional neurons with active synaptic protein expression, we identify substantially fewer neurons than most previous estimates in the *Drosophila* larval CNS, and in addition, substantially more glia. We also discover a previously unsuspected sexual dimorphism in the numbers of both cell types at larval stages. The generation of whole CNS point clouds from our data enabled us to apply the tools of topological data analysis to summarize the CNS in terms of multi-scale topological structures. Utilization of these topological summaries in a support vector machine also supports that sexual dimorphism extends to deeper features of CNS organisation. Finally, we applied our pipeline to quantitate the whole CNS expression frequency of the *Drosophila* family of voltage-gated potassium channels which revealed divergent channel expression frequency throughout the CNS. We envision that our method can be employed to allow precise quantitation of gene expression characteristics of the constituent cells of the brain and potentially other intact whole organs in a format suitable for sophisticated downstream analysis.

## RESULTS

### Genetic and imaging tools designed to facilitate automated whole CNS cellular quantitation

To establish a robust quantitative method to measure gene expression frequency and quantify the cell numbers that contribute to *Drosophila* larval CNS function, we sought to develop a pipeline utilising genetic reporters designed to expediate automated neuron and glia quantitation from three-dimensional intact organ images. While membrane associated reporters are generally employed to label *Drosophila* neurons (Pfeiffer et al., 2008; Jenett et al., 2012; Ravenscroft et al., 2020), we generated UAS-driven (Brand and Perrimon, 1993; Wang et al., 2012) fluorescent reporters fused to Histone proteins (Sherer et al., 2020) to target fluorescence only to the nucleus, in order to facilitate subsequent automated segmentation and counting. Through empirical selection of transgene genomic integration sites, we established a set of reporter lines that produced a strong and specific punctate nucleus signal when expression is induced, with little to no unwanted background expression. We then developed a procedure to capture the entire micro-dissected larval CNS volume by light sheet microscopy at multiple angles and with high resolution, imaging only animals within the ∼two-hour developmental time window of the wandering third instar larval stage (Ainsley et al., 2008). These multi-view datasets were then processed to register, fuse, and deconvolve the entire larval CNS volume. The volume was then segmented and cell numbers automatically quantified (Figure 1a-d).

**Figure 1.**
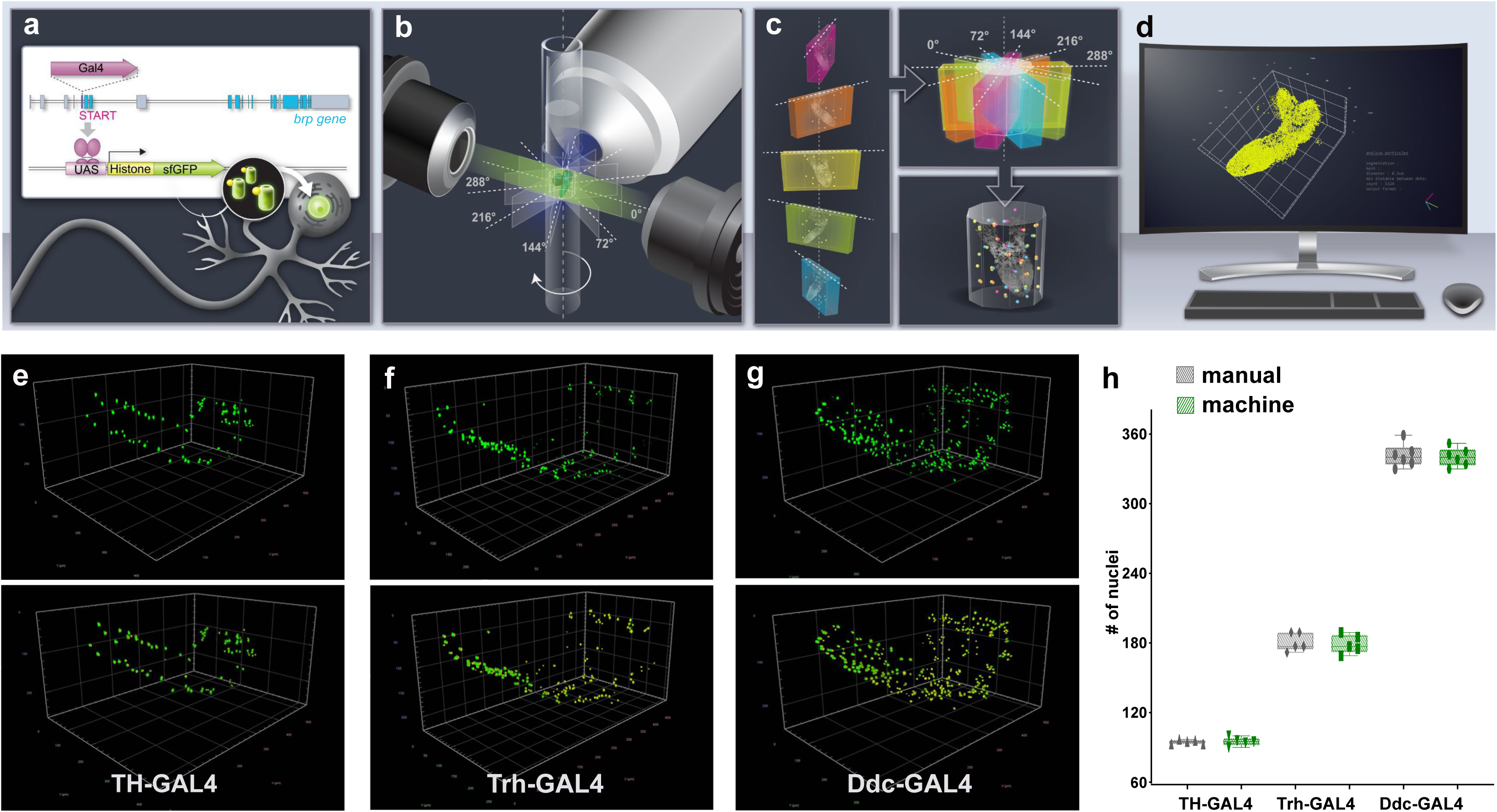
Intact whole CNS quantitation pipeline schematic and validation. (**a-d**) Illustration of intact whole CNS genetic, imaging and computational pipeline. (**a**) Genetic reagents: GAL4 is introduced to the exonic sequences of genes encoding synaptic proteins (e.g. *brp*) to capture their expression pattern with high fidelity. GAL4 expression regulates the production of UAS fluorescent-histone reporters which target to the nucleus of cells producing a punctate signal. (**b**) Imaging: The intact CNS is imaged at high resolution using light-sheet microscopy. Images are captured at 5 different angles with 72-degree intervals. (**c**) Assembly: multiview light sheet images are registered, fused and deconvolved. (**d**)Quantitation: volume is segmented, nucleus number and relative position is measured. Three dimensional co-ordinates of the geometric center of every nucleus can be calculated to produce a point cloud of nuclei positions.(**e-h**) Pipeline validation. Three dimensional images before (above) and after (below) segmentation panels of (**e**) dopaminergic [TH-GAL4] neurons, (**f**) serotonergic neurons [Trh-GAL4] and (**g**) dopa decarboxylase expressing [Ddc-GAL4] neurons. (**h**) Manual or automated quantification of nuclei numbers in these volumes are similar. Scale squares in **e** and **g** is 100μm and in **f** is 50μm. n= 6 for Ddc-GAL4, Figure 1-source data 1) between automated and manual measurements of these neuronal subtypes establishing confidence in the procedure.

To evaluate the reliability of the procedure, we began by comparing automated counts of distinct neuronal subtypes with manual counting. We separately labelled all dopaminergic neurons (Figure 1e) [TH-GAL4](Friggi-Grelin et al., 2003a; Mao and Davis, 2009), serotonergic neurons (Figure 1f)[Trh-GAL4] (Alekseyenko et al., 2010), and neurons that produce both types of neurotransmitter (Figure 1g )[Ddc-GAL4] (Lundell and Hirsh, 1994) in the larval CNS. Quantification revealed a high level of concordance (Figure 1h, +/-0.21%, n= 5 for TH-GAL4, +/-1%, n= 5 for Trh-GAL4, +/-0.38%,

### Number of neurons and glia in the female larval CNS

Encouraged by our neuronal subset quantitation results, we next sought to generate GAL4 lines for genes likely to be expressed only in active larval neurons with synaptic connections but not by neuroblasts or by immature neurons (Figure 2-figure supplement 1). We biased towards generating GAL4 insertions within endogenous genomic loci in order to reproduce endogenous patterns of gene expression with high fidelity.

Bruchpilot (Brp) is a critical presynaptic active zone component widely used to label *Drosophila* synapses, including for large-scale circuit analyses (Wagh et al., 2006). We employed CRISPR/Cas9 genome editing to insert GAL4 within exon 2 of the *brp* gene, utilising a T2A self-cleaving peptide sequence (Diao et al., 2015) to efficiently release GAL4. While this exonic insertion generated a hypomorphic allele of *brp* (data not shown) when homozygous, the line was employed in heterozygotes to capture Brp protein expression with high fidelity. To complement this line, we used the Trojan/MiMIC technique (Diao et al., 2015), to generate a GAL4 insertion in the *syt1* gene, which encodes Synaptotagmin 1 (Littleton et al., 1994), the fast calcium sensor for synaptic neurotransmitter release (Quiñones-Frías and Littleton, 2021). Lastly, we examined a transgenic line where the promoter of *nsyb* (*neuronal synaptobrevin*) (Deitcher et al., 1998), which encodes an essential presynaptic vSNARE (Südhof and Rothman, 2009), is used to control GAL4 expression (Aso et al., 2014). All three lines were expressed in a similar pattern, labelling a substantial fraction but not all of the total cells in the larval CNS (Figure 2a-c). These lines contrasted with the widely used elav-GAL4 (Lin and Goodman, 1994), which was expressed in larval neurons, but also apparently in some immature neurons and potentially in some glia as well (Berger et al., 2007) (Figure 1-figure supplement 1). To characterise our lines, we examined their expression throughout development beginning with embryogenesis. We detected no expression from any of the three lines prior to embryonic stage 16 (Figure 2-figure supplement 2a). However, beginning at stage 17 of embryonic development, when synaptic activity begins (Baines and Bate, 1998), all three lines displayed expression in both the peripheral and central nervous system (Figure 2-figure supplement 2). We also examined if these lines were expressed in neural stem cells during larval stages by co-labelling the larval CNS with the transcription factor Deadpan, a neuroblast marker (Bier et al., 1992). We found that labelling by all three lines did not overlap with Deadpan expression (Figure 2-figure supplement 2), suggesting these lines are not expressed in neuroblasts. We also examined expression all three lines in the adult brain and, as in the larval CNS, observed labelling of a large fraction but not all of the total cells in the adult brain (Figure 2-figure supplement 2). Lastly, to ensure that the cells labelled by our lines were exclusively neurons, we compared their expression to that of glial cells labelled by glial specific transcription factor Repo (Xiong et al., 1994; Lin and Potter, 2016) using independent and mutually exclusive QF2 dependent labelling. We found complete exclusion of cells labelled by Brp, Syt and nsyb from cells labelled by Repo (Figure 2d), consistent with the Brp, Syt1 and nsyb Gal4 lines labelling only neurons that express synaptic protein genes but not glial cells.

**Figure 2.**
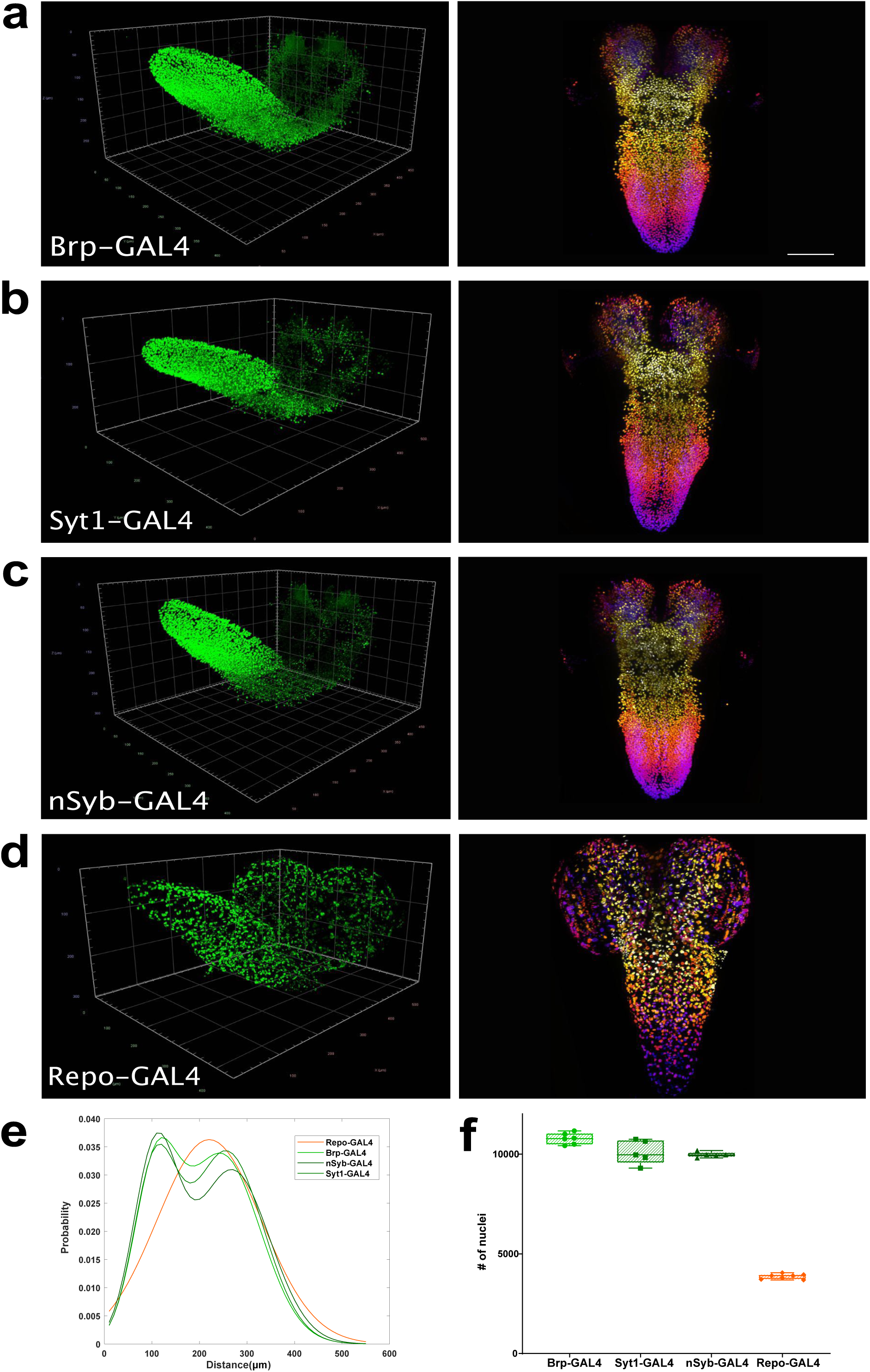
Whole CNS quantitation of neurons and glia in the female larval brain. (**a-d)** Multiview deconvolved images (left) and z-stack projections (right) [colours represent z position] of the CNS of (**a**) Brp-GAL4, (**b**) Syt1-GAL4, (**c**) nSyb-GAL4 and (**d**) Repo-GAL4. (**e**) Distribution of inter-nuclei distances for each line. (**f**) Quantification of the number of labelled nuclei in each line. **a-d**: left; scale squares **a** and **c** = 50μm, **b** and **d** = 100μm; right images identical magnification, scale bar = 100μm.

To further compare these lines, beginning with the CNS of female animals, we calculated three dimensional coordinates for the geometric center of all nuclei labelled in the Brp, Syt1 and nSyb GAL4 lines to generate point cloud mathematical objects and compared them to point clouds of glial nuclei labelled by the Repo GAL4 line. We then plotted and compared the distributions of inter-nuclei distances in these lines. Using this measurement, we found that inter-nuclei distance of glial cell nuclei exhibited a unimodal distribution (Figure 2e). In contrast, all three neuronal lines exhibited a bimodal distribution of inter-nuclei distances (Figure 2e). We thus observed two patterns of labelled nuclei, one shared among neuronal lines and the other distinct for glia (Figure 2e), again consistent with these lines labelling different cell types.

We next counted the number of nuclei labelled by these neuronal and glial lines, again beginning with females (Figure 2f). We found that the CNS labelled by Brp-Gal4 had 10776(+/− 2.65%, n= 6) neurons, Syt1-Gal4 had 10097(+/− 5.96%, n= 5) neurons and nSyb-Gal4 had 9971(+/− 1.35%, n= 5) neurons (Figure 2f). We tested the statistical difference in the numbers of neurons labelled by these lines and found that while nSyb-GAL4 and Syt1-GAL4 were not statistically different from each other, Brp-GAL4 did label significantly more neurons than either Syt1 or nSyb (Brp-GAL4 vs Syt1-GAL4 + 6.72%, p=0.03, Brp-GAL4 vs nSyb-GAL4 + 8.07% p=0.01). Averaging across the lines, we found that the female third instar larval CNS had 10312 +/− 5.03%, n=16 neurons (Figure 2-source data 1). To ensure that our method did not introduce bias in dense data sets, we also manually counted a Brp-GAL4 labelled CNS and compared it to the automated count. Similar to our experiments with sparse neuronal labelling, we found good agreement between manual and automated quantification with a difference of just 14 neurons (9430 nuclei manual vs 9444 nuclei automated for this individual CNS).

We next counted the number of glia labelled by the Repo GAL4 line (Figure 2d,f). We measured 3860 +/− 3.37%, n=7 glia in the female CNS (Figure 2-source data 1). This amounted to 37% of the number of neurons, far more than the previously estimated (Meinertzhagen, 2018; Raji and Potter, 2021). In sum, we found that the female *Drosophila* larval CNS had 10312 neurons, fewer than most previous predictions and several fold more glia that previously thought.

### Males have fewer neurons and more glia than females

We next carried out a similar analysis on the CNS of male larvae (Figure 3a-c). We found that Brp-GAL4 labelled 9888 (+/-3.15%, n= 5) neurons, Syt1-GAL4 labelled 9012 (+/-3.8%, n= 5) neurons, and nSyb-GAL4 labelled 9286 (+/− 5.38%, n= 5) neurons in male larvae (Figure 3e, Figure 3-source data 1). In males, Brp-GAL4 did not label significantly more neurons than nSyb-GAL4 but did label more than Syt1-GAL4 (Brp-GAL4 vs Syt1-GAL4 + 876, p=0.01), while the number of neurons labelled by nSyb-GAL4 was not significantly different from Syt1-GAL4, as we had found in females. Averaging across the lines we found that the male third instar larval CNS had 9396 +/− 5.59%, n= 15 neurons, significantly fewer than those of females (−9.75%, P<0.0001). This difference was also consistent within individual genotypes with Brp-GAL4 labelling (−8.98%, P=0.0008), Syt1-GAL4 labelling (−12.04%, P=0.008) and nSyb-GAL4 labelling (−7.38%, P=0.0182) less neurons in males than in females.

**Figure 3.**
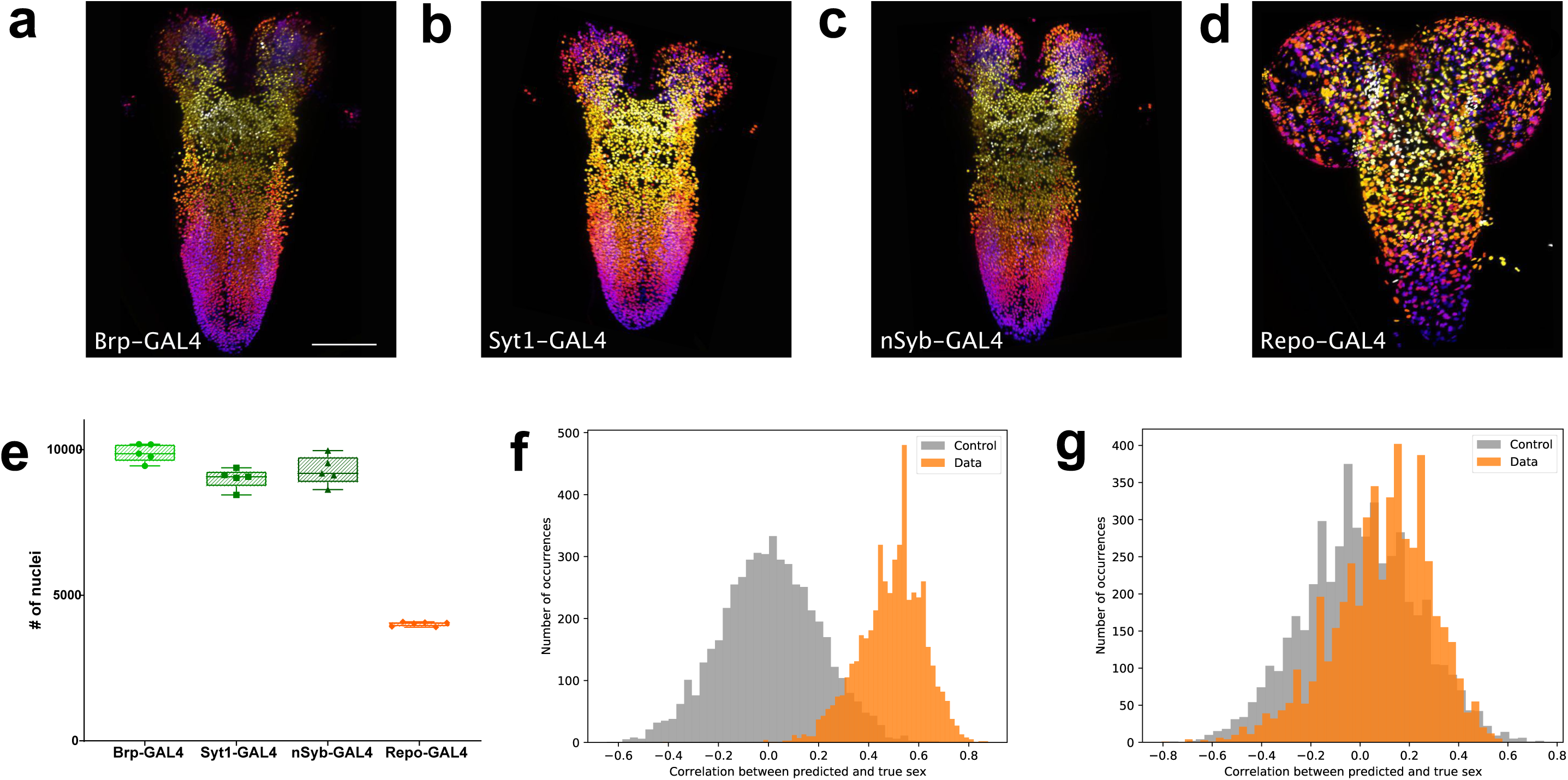
Quantitation of neurons and glia in the male larval CNS and topological comparison of sex differences. (**a-d**) Example z-stack projections [colours represent z position] of male larval CNS of (**a**) Brp-GAL4, (**b**) Syt1-GAL4, (**c**) nSyb-GAL4 and (**d**) Repo-GAL4.(**e**) Quantification of the number of labelled nuclei in each line. (**f**) The distribution of correlations between the ground truth and the prediction made by the SVM using topological features is indicative of sexual dimorphism of the higher order structure of neuron point clouds (**g**) Simpler point cloud features such as properties of the distributions of inter-nuclei distances are not indicative of this. **a**-**d**: identical magnification, scale bar =100μm

We also counted the number of glia labelled by Repo-GAL4 in males (Figure 3d, e). We found that males had 4015 glia, again far more than previous estimates (Figure 3-source data 1). The number of glia in the male larval CNS was also significantly more than in females (+3.86%, P=0.0284). In summary, male *Drosophila* larva have significantly fewer CNS neurons than females but significantly more glia.

### Topological analysis detects significant structural differences between males and females

We next wished to determine whether the differences between the point clouds derived from the positions of neuronal nuclei of males and females went beyond simple numerics. To do this we applied the tools of topological data analysis (Rabadan and Blumberg, 2019; Chazal and Michel, 2021) to summarize the CNS in terms of multi-scale topological structures (Expert et al., 2019). These topological summaries, the construction of which is described in the methods, can be thought of as multi-scale descriptions of the shape of the data set. Topological summaries, which can be compared by standard methods despite the lack of common reference points, could then be used as the classification features in a support vector machine (SVM). Since the total number of point clouds was relatively small for this type of analysis, we down-sampled each whole CNS point cloud randomly to 8000 points 100 times, producing a total of 3100 point clouds, for each of which we then computed a certain topological summary, called the *degree-1 persistence diagram of its alpha complex* (Edelsbrunner and Mücke, 1994).

After fixing the necessary hyperparameters, sex classification experiments were run across 5000 random train/test splits of the topological summaries. In each split, the summaries derived from subsamplings of a single point cloud (CNS) were either all in the training set or all in the testing set, to avoid leaking information. Each time, the SVM was trained once with the animal’s true sex as the target class and once with a randomly assigned sex as the target, as a control. We then computed the Pearson correlation between the classifier’s output on the testing set and the true (respectively randomized) sex of the animal.

The 5000 splits were used to produce 5000 correlations with the true sex and 5000 correlations with a randomly assigned sex. The distribution of these correlations (Figure 3f), exhibiting clearly that the SVM is able to extract the sex of the animal reliably: only about 1.9% of the splits result in a higher correlation in the control set than in the true data. Moreover, repeating the procedure with simpler point cloud features, like properties of the distributions of inter-nuclei distances, did not produce a significant signal (Figure 3g). Thus, the pattern, which seems hard to describe concisely, is not revealed through simpler descriptors of the neuron configurations, leading us to suspect CNS sexual dimorphism extends to deeper features of organisation that are both subtle and widely distributed. These results, in addition to the differences in total cell numbers, support sexual dimorphism of the male and female CNS at the larval stage.

### Potassium channel family members have different densities in the CNS

Having established a baseline of total numbers of neurons in the larval CNS, we next sought to deploy the quantification pipeline to measure the expression frequency of key neuronal function genes throughout the CNS. We chose to examine the family of voltage-gated potassium channels, which are essential for many aspects of neuronal function and for which *Drosophila* studies defined the founding members (McCormack, 2003). We generated GAL4 insertions in the Shaker (Sh) [Kv1 family], Shab (Sb) [Kv 2 family], Shaw (Sw) [Kv3 family] and Shal (Sl) [Kv4 family](McCormack, 2003) genes using the Trojan/Mimic technique (Diao et al., 2015). As the Sh gene is x-linked, we carried out our quantitation analysis in the male CNS only to avoid potential gene dosage effects. To determine whether our GAL4 reporter lines had patterns of expression consistent with the known properties of these channels, we examined the expression of all four lines in motor neurons, where functional activity for Shaker, Shab, Shaw and Shal has previously been demonstrated by electrophysiological measurements (Covarrubias et al., 1991; Ryglewski and Duch, 2009). We found that the GAL4 reporters for all 4 channels were expressed as expected in motor neurons (Figure 4-figure supplement 1), consistent with accurate reproduction of the established expression of these proteins.

We next examined the expression frequency of these genes in the entire CNS (Figure 4a-d). We found that Shaker and Shal were expressed in large numbers of neurons 8204 +/− 5.67% and 8261 +/− 3.1%, though significantly less (−12.7 % and −12.1% p<0.0001) than the average number of all male neurons (Figure 4 a,b,e, Figure 4-source data 1). In contrast, Shab (3057+/− 8.21% n=10) and Shaw (1737 +/− 4.3% n=11) were expressed in smaller numbers of neurons (Figure 4c-e), with expression observed in only 32.5% or 18.5% of total male neurons respectively, suggesting more discrete functions within CNS neurons and contrasting with the collective expression of all four genes within motor neurons (Figure 4-figure supplement 1). In particular, Shab and Shaw had very reduced expression in the brain lobes of larva (Figure 4c,d) compared with Shaker and Shal (Figure 4a,b). These results establish that our genetic-imaging pipeline can enable quantitation of the expression frequency of families of genes essential for neuronal properties across the entire CNS.

**Figure 4.**
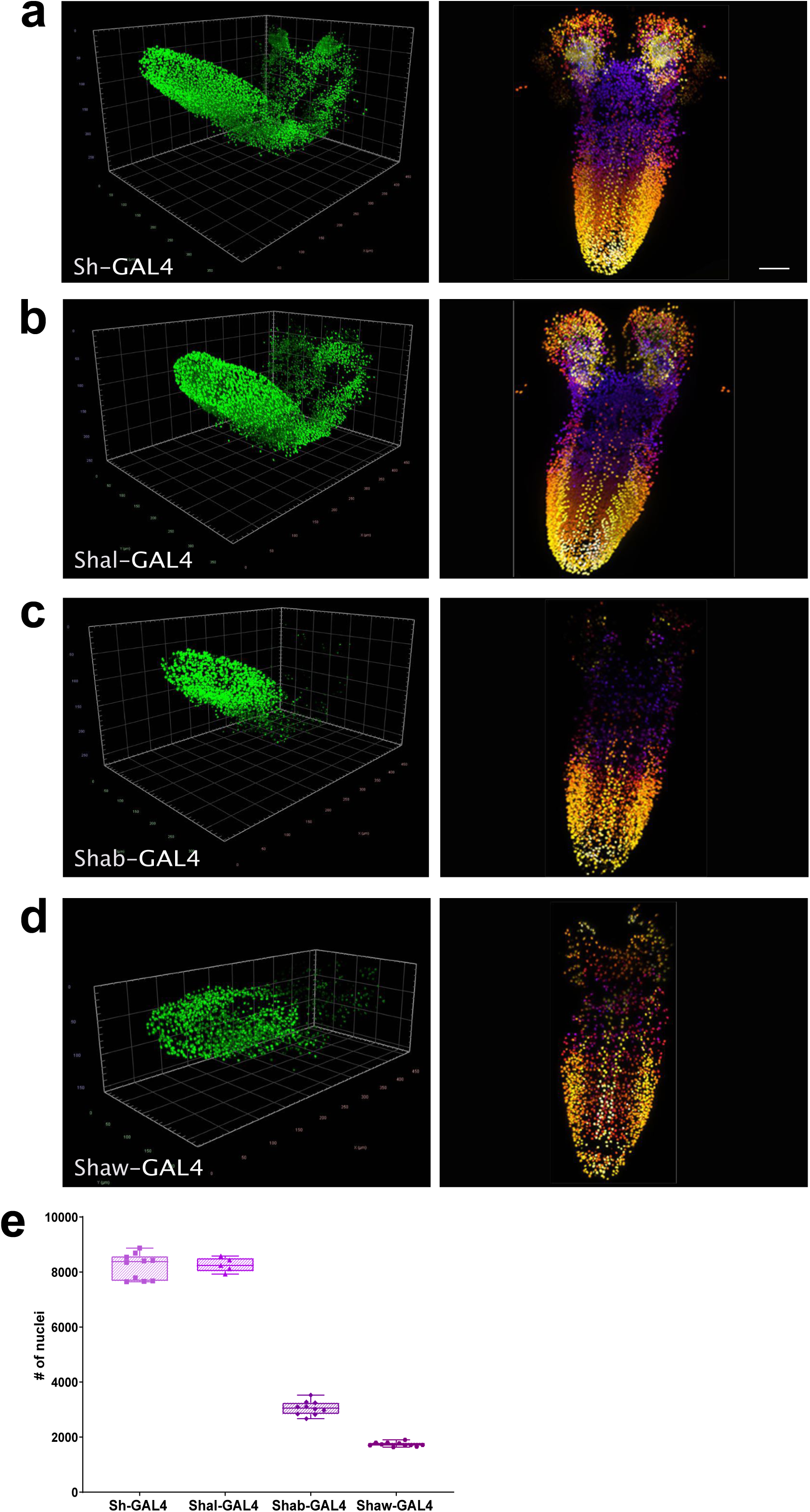
Quantitation of the number of neurons expressing voltage-gated potassium channels. (**a-d**) Multiview deconvolved images (left) and z-stack projections (right) [colours represent z position] of potassium channel family members: (**a**) Sh-GAL4, (**b**) Shal-GAL4, (**c**) Shab-GAL4 and (**d**) Shaw-GAL4. (**e**) Quantification of the number of labelled nuclei in each line. **a-d**: left, scale squares = 50μm, right, identical magnification, scale bar = 50μm.

## DISCUSSION

Establishing the number and identity of cells in the CNS is a foundational metric upon which to construct molecular, developmental, connectomic and evolutionary atlases of central nervous systems across species (Lent et al., 2012; Devor et al., 2013). Here, we develop and deploy a methodological pipeline to label discrete cell types in the intact *Drosophila* CNS, with genetic reporters designed to facilitate the subsequent segmentation and automated quantification of cell types, in addition to capturing positional coordinates of relative nucleus position throughout the organ. Using this toolset, we find fewer active neurons, as defined by expression of synaptic protein genes, in the *Drosophila* larval CNS than the majority of previous predictions and substantially more glia. We also discover previously unsuspected differences in both neuron and glial density and CNS topology at the larval stage, when external sex organs are absent, with females possessing both more neurons and but fewer glia than males. Topological analysis of the point cloud derived from neuronal nucleus position, which detects potentially subtle and complex geometric structure in the data, also strongly supports the existence differences between males and females. In addition, deploying these tools, we find that while all members of the *Drosophila* voltage-gated potassium channel family are expressed in motor neurons, consistent with prior mutant analyses, the Kv2 channel Shab and Kv3 channel Shaw are expressed in a much smaller number of neurons in the CNS than the Kv1 channel Shaker and the Kv4 channel Shal, suggesting conclusions drawn about the coordinated activity of these channels from studies of motor neurons may not be broadly applicable across the CNS, where the genes encoding these channels are frequently not co-expressed.

A number of semi-quantitative methods have been employed to estimate the number of neurons in the brains of humans and model organisms, including *Drosophila* (Lent et al., 2012; Keller et al., 2018). For example, the number of neurons or other cells in the brain has been estimated using stereological counting of sub-regions. A major limitation of this approach is the assumption of homogenous cell density across the organ or within subregions, which is not supported by the high variability of counts even between samples of similar regions, and thus likely introduces large errors (von Bartheld et al., 2016; Keller et al., 2018). Rough extrapolation of neuronal counts of electron microscope volumes of the regions of the *Drosophila* larval CNS had suggested an estimate of ∼15,000 neurons (Meinertzhagen, 2018; Eschbach and Zlatic, 2020). An alternate approach is isotropic fractionation, where all cells in large regions or the entire CNS are dissociated to produce a homogeneous single-cell suspension. Nuclei in the suspension can then be labelled by immunohistochemistry and cells in a subvolume counted in a Neubauer chamber to estimate the total number of cells present. Limitations of the approach include the necessity to ensure complete dissociation of cells while avoiding tissue loss, the requirement for homogenous antibody labelling, and highly accurate dilution (Deniz et al., 2018). This approach has recently been used to estimate the total number of neurons and glia in the adult *Drosophila* brain and suggested a number of 199,000 neurons (Raji and Potter, 2021), twice prior estimates (Scheffer and Meinertzhagen, 2019; Allen et al., 2020). In contrast to our results in the larval CNS, this study found no significant differences in the number of neurons between the sexes and also found that ‘non-neuronal’ cells, which should include glia, accounted for less than 9% of the total cells counted. In addition to the inherent inaccuracy of the isotropic fractionation technique, which the authors both observed and acknowledge (Raji and Potter, 2021), their use of anti-Elav antibody labelling, which can label some glia in addition to neurons (Berger et al., 2007), or perhaps differences in life stage, may explain some of the discrepancies between our results.

An unpredicted result from our whole CNS neuron quantitation was substantial differences in neuron and glial numbers between the sexes in larva. In adult *Drosophila*, sexually dimorphic neural circuitry has been observed in olfactory system (Kimura et al., 2005), and human females have also been reported to have more olfactory bulb neurons and glia than males (Oliveira-Pinto et al., 2014). While sex-specific behavioural differences are obvious in adult *Drosophila* (Jazin and Cahill, 2010), few sexually dimorphic behavioural differences have been reported in larva (Aleman-Meza et al., 2015). However male and female larva do differ in nutritional preference (Rodrigues et al., 2015; Davies et al., 2018), which could potentially account for some aspects of the dimorphism we observe. In addition to differences in total cell numbers, our topological methods, which take into account multi-scale structure, suggests that differences in CNS structure between the sexes is both subtle (in the mathematical sense) and non-localised in nature, and indeed are not observable with simpler analysis methods of CNS organisation.

In addition to enabling precise counting of genetically labelled cells, our method allows the relative measurement of discrete cell types or gene expression frequencies throughout the CNS. For example, the relative frequency of glial cells to neurons in the human brain has been long been debated (von Bartheld et al., 2016) and in the adult *Drosophila* brain it has been suggested there are 0.1 glial per neuron (Kremer et al., 2017; Scheffer and Meinertzhagen, 2019; Raji and Potter, 2021). In the larval *Drosophila* CNS, we found closer to 0.4 glial cells per neuron on average, more similar to the glial-neuron ratios reported for rodents or rabbits (Verkhratsky and Butt, 2018). An important potential caveat however is that the large relative ratio of glia we observe in the third instar larva could conceivably be glia produced in advance of adult CNS development. As adult specific neuron numbers expand during pupation, the relative ratio of glia could potentially decline. Additional glial-neuron ratio measurements in the adult CNS will be required to examine this possibility.

Our approach may also allow assignment of potential functional classes of neurons. For example, from our examination of voltage-gated potassium channel family gene expression, all are collectively expressed in motor neurons, however the Shab and Shaw genes have more discrete expression patterns in other CNS neuron classes, potentially imbuing these neurons with unique functional characteristics (Chow and Leung, 2020). Future multiplexing of binary genetic expression systems and reporters (Simpson, 2009; del Valle Rodríguez et al., 2011; Diao et al., 2015) should enable neurons or glia to be further quantitively sub-classified by gene expression features throughout the entire intact central nervous system.

## MATERIALS AND METHODS

### Drosophila stocks

The following stocks were employed - y[1] w[67c23]; Mi{PT-GFSTF.0}Syt1[MI02197-GFSTF.0]/CyO (BDSC#59788)(Venken et al., 2011), y[1] w[*] Mi{y[+mDint2]=MIC}Sh[MI10885] (BDSC#56260), y[1] w[*];Mi{y[+mDint2]=MIC}Shal[MI10881] (BDSC#56089)(Venken et al., 2011), y[1] w[*]; Mi{y[+mDint2]=MIC} Shab[MI00848] (BDSC#34115)(Venken et al., 2011), nSyb-GAL4(R57C10)(Pfeiffer et al., 2008), repo-GAL4 (BDSC#7415)(Sepp et al., 2001), repo-QF2 (BDSC#66477)(Lin and Potter, 2016), Shaw-TrojanGAL4 (BDSC#60325)(Venken et al., 2011; Li-Kroeger et al., 2018), Ddc-GAL4(BDSC#7009)(Feany and Bender, 2000), TH-GAL4(BDSC#8848)(Friggi-Grelin et al., 2003b), Trh-GAL4(BDSC#38389)(Alekseyenko et al., 2010), UAS_H2A-GFP(Sherer et al., 2020), QUAS_H2B-mCherry(Sherer et al., 2020), Brp-GAL4 (this manuscript), UAS_H2A::GFP-T2A-mKok::Caax (this manuscript). All lines were raised on standard media at 25°C, 50%RH.

### Generation of Brp-GAL4 exon 2 insertion line

A GAL4.2 sequence was inserted in genome, immediately after the start codon of the Brp-RD isoform using CRISPR based gene editing employing the following constructs. *Brp gRNA pCDF3:* Two gRNA sequences targeting each side of the insertion location in exon 2 of *brp*, were selected using the FlyCRISPR algorithm (http://flycrispr.molbio.wisc.edu/), consisting of 20 nucleotides each (PAM excluded), and predicted to have minimal off-targets. Each individual 20-nucleotide gRNA sequence were inserted into pCFD3 plasmid (Addgene #49410) using the KLD enzyme mix (New England Biolabs). *Brp-GAL4 insertion construct*: The 7 following PCR amplified fragments were assembled using HIFI technology - (1) 1198bp Homology arm covering 5’UTR until 5’ target site; (2) the region between 5’ target site and the start codon were amplified from drosophila nos-cas9 (attp2) genomic DNA (a modified Pam sequence was inserted using overlapping primers); (3) Linker-T2A-GAL4.2 sequence was amplified from pBID-DSCP-G-GAL4 (Wang et al., 2012) (the linker-T2A sequence was added upstream of the forward primer); (4) P10-3’UTR was amplified from pJFRC81-10XUAS-IVS-Syn21-GFP-p10 (Addgene 36432); (5) 3xP3-Hsp70pro-dsRed2-SV40polyA selection cassette, flanked by two LoxP sites, was amplified from pHD-sfGFP Scareless dsRed (Addgene 80811); (6) The region covering the end of DsRed cassette until 3’ target site and (7) the 1079bp Homology arm 2 covering from the 3’ target site to exon 2, were amplified from Drosophila nos-cas9 (attp2) genomic DNA. Full length assembly was topo cloned in zero-blunt end pCR4 vector (Invitrogen), all constructs have been verified by sequencing (Microsynth AG, Switzerland) and injections were carried out into a nos-cas9 [attp2] strain (Ren et al., 2013). Correct insertion of GAL4 was verified by genome sequencing. All primer sequences are included in the Key Reagents table.

### Construction of UAS_H2A::GFP-T2A-mKok::Caax

PCR amplifications were performed using Platinium Superfi polymerase (Invitrogen). The three PCR fragments were assembled together using Hifi technology (Invitrogen) - (1) Histone2A (H2A) cDNA was amplified from *pDESTP10 LexO-H2A-GFP* template [Gift from Steve Stowers] with a synthetic 5’UTR sequence (syn21) added upstream to H2A on the forward primer; (2) sfGFP was amplified from template pHD-sfGFP Scareless dsRed (Addgene 80811) and (3) mKok amplified from pCS2+ ChMermaid S188 (Addgene 53617) with the CAAX membrane tag sequence (Sutcliffe et al., 2017) added at the 3’ end of the protein using the reverse primer. A *Thosea asigna* virus 2A(T2A) self-cleaving peptide sequence (Diao et al., 2015), was inserted between sfGFP and mKok, using sfGFP reverse and mKok forward overlapping primers. The full length assembly was TOPO cloned into pCR8GW-TOPO vector (Invitrogen) generating pCR8GW-H2A::GFP-T2A-mKok::Caax. The insert, H2A::GFP-T2A-mKok::Caax was , then, transferred to pBID_UASC_G destination vector (Wang et al., 2012) using LR II clonase kit (Invitrogen) to generate pBID_UAS-H2A::GFP-T2A-mKok::Caax. The transgene was generated by injection into the JK66B landing site. All primer sequences are included in the Key Reagents table.

### Generation of novel Trojan GAL4 lines

The MiMIC lines generated by the group of Hugo Bellen(Venken et al., 2011) were acquired from the Bloomington Stock Center. Conversion of Mimic lines to Trojan GAL4 lines lines was performed as described previously (Diao et al., 2015).

### Larval CNS preparation and image acquisition

Wandering 3^rd^ instar larvae were dissected in 1x PBS (Mediatech) and fixed with 4% formaldehyde (Sigma-Aldrich) for 20 mins. 1x PBS were added to remove the fixative, and then the CNS was dissected(Hafer and Schedl, 2006) and rinsed with 1xPBS with 4% Triton-X 100 for 2 days at 4°C. After rinses, the CNS was embedded in 1% low melting temperature agarose (Peq gold) mixed with 200nm red fluorescent beads (1:50000), then introduced into a glass capillary and positioned well separated from each other. After solidification of the agarose, the capillary was mounted to sample holder, transferred to a Zeiss Lightsheet Z.1 microscope and the samples were extruded from the capillary for imaging. CNS images were acquired with a 20x/1.0 Apochromat immersion detection objective and two 10x/0.2 illumination objectives at 5 different views, with 1µm z-intervals. Voxel resolution was 0.317um.

### Image processing and data analysis

Collected multiview datasets were registered and fused with the Fiji Multiview Reconstruction plugin(Preibisch et al., 2010; Schindelin et al., 2012). Image datasets after Multi-view deconvolution were analyzed with Vision4D 3.0.0 (Arivis AG). A curvature flow filter was first used to denoise the image dataset. Subsequently, a Blob Finder algorithm(Najman and Couprie, 2003) was applied to detect and segment bright rounded 3D sphere-like structures in the images with 4.5µm set as the diameter. Segmented objects with volume less than 15µm^3^ were removed from analysis by segmentation filter to avoid unspecific signals. Subsequently, the number of nuclei and the x, y, z coordinates of the center geometry of each nucleus were output from Vision4D. Where manual counting was employed (Figure 1 and a randomly selected Brp-GAL4 labelled CNS), Vision4D was used to visualize and iteratively proceed through and manually annotate the dataset. Example whole CNS datasets where functional neurons or glia are labelled are available (Jiao and McCabe, 2021a, 2021b). In 2D representations, Z position is indicated by colour coding using the scheme below.

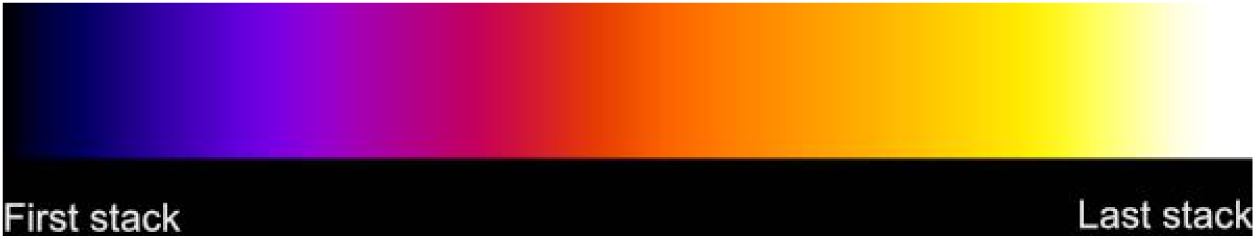

### Mathematical analysis

The topological summaries methods employed have previously been introduced (Edelsbrunner and Harer, 2010; Ghrist, 2014; Rabadan and Blumberg, 2019; Chazal and Michel, 2021). For a motivating example of the principles underlying topological summaries, one could think of pearls forming a necklace. Topological summaries express the global structure of the necklace formed by the relationships between the positions of the individual pearls, but are invariant under translation and rotation of the necklace overall. Two such necklaces have topological summaries that are comparable even if the pearls in one have no relationship to the pearls in the other. It is the global structures – such as its circular shape on a large scale, or bulges on a smaller scale – formed by the relationships of the individual pearls of each one that matter.

We trained a machine learning classifier, specifically an SVM (support vector machine), on the CNS nuclei positions, in order to evaluate its power in determining characteristics of the animal from which it was derived. The data encompassed all point clouds generated from all CNS lines (Brp-Gal4, Syt-Gal4, nSyb-Gal4, Repo-Gal4). Correlation significance (classification power) is determined by comparing the performance of the SVM on the actual classification task to one wherein each larva is randomly assigned a class.

Mathematically speaking, the nuclei positions from a single CNS form a point cloud, a finite set of points in R^3^. A possible, naive approach to SVM feature selection for point clouds would be to consider the mean, variance, or other modes of the distribution of pairwise distances within the cloud. These real-valued features could then be passed through, for example, radial basis function kernels for use in SVMs. We focused on very different kind of features, namely ones obtained from the topology of the point clouds. When the point cloud is of low dimension, such as the three-dimensional point clouds arising from nuclei position data, the following approach is relevant. Let X be a finite point cloud in R^3^. For any r ≥ 0, we let X_r_ denote the same point cloud, but with each point replaced by a ball of radius r. As r increases, the sequence formed by the X_r_ expresses different topological features of X. By topological features, we here mean the presence or absence of multiple connected components, unfilled loops, and unfilled cavities.

The figure below illustrates this process in the case of a synthetic 2-dimensional point cloud, but the idea extends to any dimension including whole CNS point clouds. When r is small, X_r_ is topologically very similar to X = X_0_, and is essentially a collection of disjoint points. When r is very large, X_r_ is topologically very similar to X_∞_, i.e., one giant, featureless blob. As the sequence X_r_ progresses through the continuum of scales between these two trivial extremes, it undergoes non-trivial topological changes: components merge, and loops form and later get filled. In higher dimensions, cavities of various dimensions likewise form and get filled in.

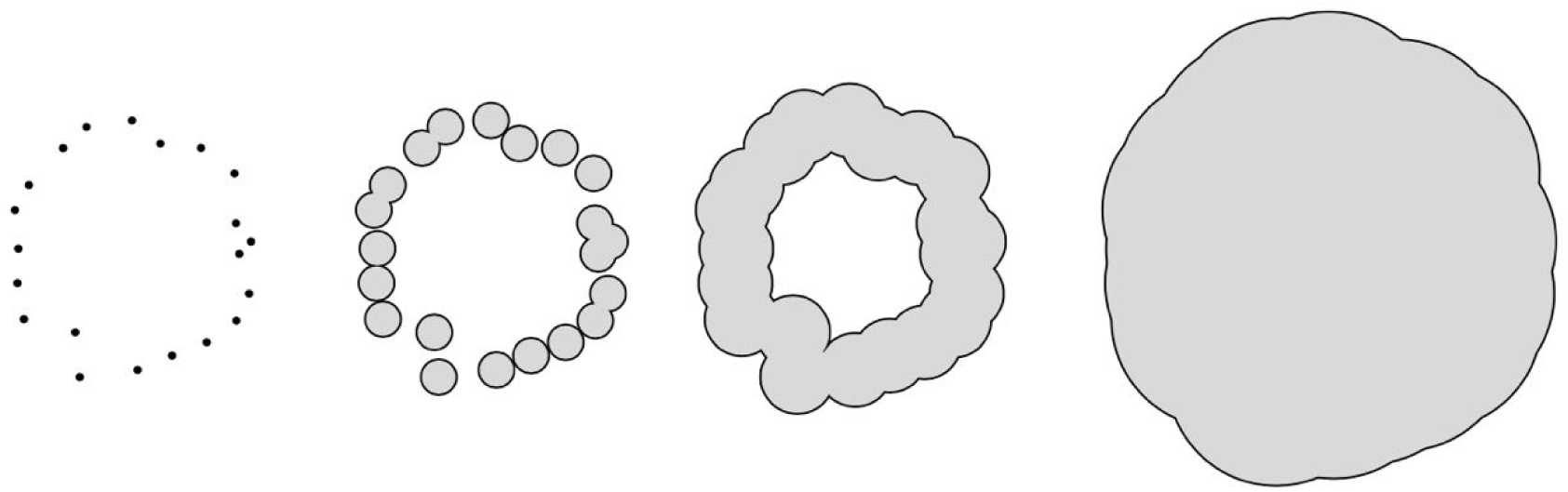

A small 2-dimensional point cloud X viewed at four different scales 0 < a < b < c, forming the filtration X = X_0_ ⊂ X_a_ ⊂ X_b_ ⊂ X_c_.

In the parlance of topological data analysis (TDA), we refer to this appearance and disappearance of topological structures as the *birth* and *death* of homology classes in various degrees. We capture the whole life cycle with a mathematical object called the *persistent homology* of the point cloud, which can be fully described by its *persistence diagram*, a planar collection of points (labelled by multiplicity), whose coordinates encode the birth and death of homological features. For the filtration in the figure above, the persistence diagram that tracks 1-dimensional features (i.e., unfilled loops) contains only a single point with coordinates (x, y). Here the first coordinate, x, is the radius at which the loop is first formed, and the second coordinate, y, is the radius at which the loop has just been filled in. In the example it is clear that a < x < b < y < c.

As multisets of points in the plane, persistence diagrams are not immediately usable as features for SVMs. One way to vectorize persistence diagrams and thus render them digestible by SVMs is to define kernels based on the diagrams, with the heat kernel (Reininghaus et al., 2015) being an oft-used candidate with nice properties. For persistence diagrams P and Q, the heat kernel can informally be defined by the inner product of two solutions of the heat equation — one with an initial condition defined by P, and the other with one defined by Q.

In this analysis in this manuscript, we calculated the persistent homology of the alpha complex (Edelsbrunner and Mücke, 1994) of the point clouds, using GUDHI(The GUDHI Editorial Board, n.d.). The heat kernels were computed using RFPKOG (Spreemann, 2021). Only the persistence diagrams in degree 1 were used. Since the number of whole CNS point clouds was relatively small, we subsampled each one randomly to 8000 points 100 times, producing a total of 3100 point clouds. This was done both in order to test the stability of the method and to ensure that the variability in the number of points in each cloud is not the source of any signal.

The hyperparameters involved, i.e., the SVM regularizer and the heat kernel bandwidth, were determined by a parameter search in the following way. Six point clouds from males and six from females were randomly selected. All 100 subsampled versions of each of these 12 constituted a training set, for a total of 1200 training point clouds. The remaining 1900 subsampled point clouds constituted the testing set. The Pearson correlation between the gender predicted by the SVM on the testing set and the ground truth was computed for each choice of hyperparameters, and a choice in a stable region with high correlation was selected: a regularization parameter C = 10 in the notation of Pedregosa *et al* (Pedregosa et al., 2011) and a bandwidth of σ = 1/100 in the notation of Reininghaus *et al* (Reininghaus et al., 2015). For the simple distance distribution features, a similar parameter selection process yielded C=10 and a radial kernel bandwidth of 10^5.

### Immunofluorescence Staining and confocal microscopy

Embryos were collected and staged at 25°C on apple agar plates supplemented with yeast paste. Standard methods were used for dechorionation, removal of the vitelline membrane and fixation(Bashaw, 2010). Embryos were stored in 100 % ethanol at −20°C before IHC labelling. Embryos were stained with mouse anti-myc^9EH10^ (1:100, DSHB), visualised with goat anti-mouse::AF488 (1:400, Jackson ImmunoResearch) together with conjugated goat anti-HRP::AF647 (1:200, Jackson ImmunoResearch) and mounted in VectorShield (Vector Laboratories). Duel colour Z-stack images of stage 15/16 and late stage 17 embryos were obtained on a CSU-W1 Confocal Scanner Unit (Yokogowa, JPN) using two prime BSI express cameras (Teledyne Photometrics). For motor neuron and adult brain preparation, larval fillets from the 3^rd^ instar larvae, or brains from adults were dissected and fixed with 4% formaldehyde (Sigma-Aldrich) for 20 mins. After fixation, samples were rinsed with 1x PBS and were washed in PBT overnight at 4°C, and then mounted in VECTASHIELD antifade mounting medium. Z-stack images were taken from Leica SP8 upright confocal microscope.

For Deadpan staining, brains from the 3^rd^ instar larvae were dissected out and fixed with 4% PFA for 20 mins. After fixation, samples were rinsed with 1x PBS, and permealized with PBT (1xPBS + 4% Triton-X 100). Antibody stainings were done in PBT + 5% normal goat serum. The dilution for chicken anti-GFP (Abcam) is 1:500, for rat anti-Deadpan (Abcam) is 1: 50. Alexa Fluor 488 anti chicken, and Alexa Fluor 594 anti-rat secondary antibodies were obtained from ThermoFisher and used at the 1:500 dilutions. Brains were mounted in VECTASHIELD antifade mounting medium, and imaged with Leica SP8.

### Statistical Analysis

Column statistics analyses were performed using GraphPad Prism 9 (GraphPad Software). For Figure 1, statistical significance was determined by unpaired t test. For Figure 2,3,4, statistical significances were determined by Ordinary one-way ANOVA, followed by a Tukey’s honestly significant difference test when multiple comparisons were required. The distribution analysis in Figure 2 were performed using matlab (MathWorks). Distances between nuclei coordinates were calculated in matlab using code available at https://doi.org/10.5281/zenodo.6574838 and plotted as a histogram of distance distribution.

## ACKNOWLEDGEMENTS

This work was supported in part using the resources and services of the BioImaging & Optics Platform (BIOP) Research Core Facility at the School of Life Sciences of EPFL and we are especially thankful for the assistance of Arne Seitz. We are grateful to Hugo Bellen, Benjamin White, Gerry Rubin, Vanessa Auld, Christopher Potter, Mel Feany, Serge Birman, Ed Kravitz, Chris Doe and Pavel Tomancak for generating Drosophila stocks, software and providing advice. Stocks obtained from the Bloomington Drosophila Stock Center (NIH P40OD018537) were used in this study.

## AUTHOR CONTRIBUTIONS

W.J, G.S, E.R, S.B, J.A, Y.S., S.S, K.H. and B.M. conceived and designed the study. W.J performed the majority of experiments and analysed the data, assisted by G.S. and Y.S. E.R. generated all novel transgenes, the Brp>GAL4 genome edit and generated illustrations. S.B. generated the Syt1 Trojan line and other useful reagents and SV imaged embryos. S.S. generated the histone fusion transgenes. W.J., G.S, K.H. and B.M. wrote the manuscript

## DATA AND MATERIALS AVAILABILITY

All genetic and molecular reagents are described in the Key Resources Table. Drosophila lines not available from public stock centers are freely available upon request – see https://www.epfl.ch/labs/mccabelab/resources/. Individual whole CNS quantitation data is available as source data files associated with each figure. Matlab code is available at https://doi.org/10.5281/zenodo.6574838. Example unprocessed whole CNS microscopy data is publicly available for neurons - doi:10.5281/zenodo.5585334 and for glia doi:10.5281/zenodo.5585358.

## Key Resources Table

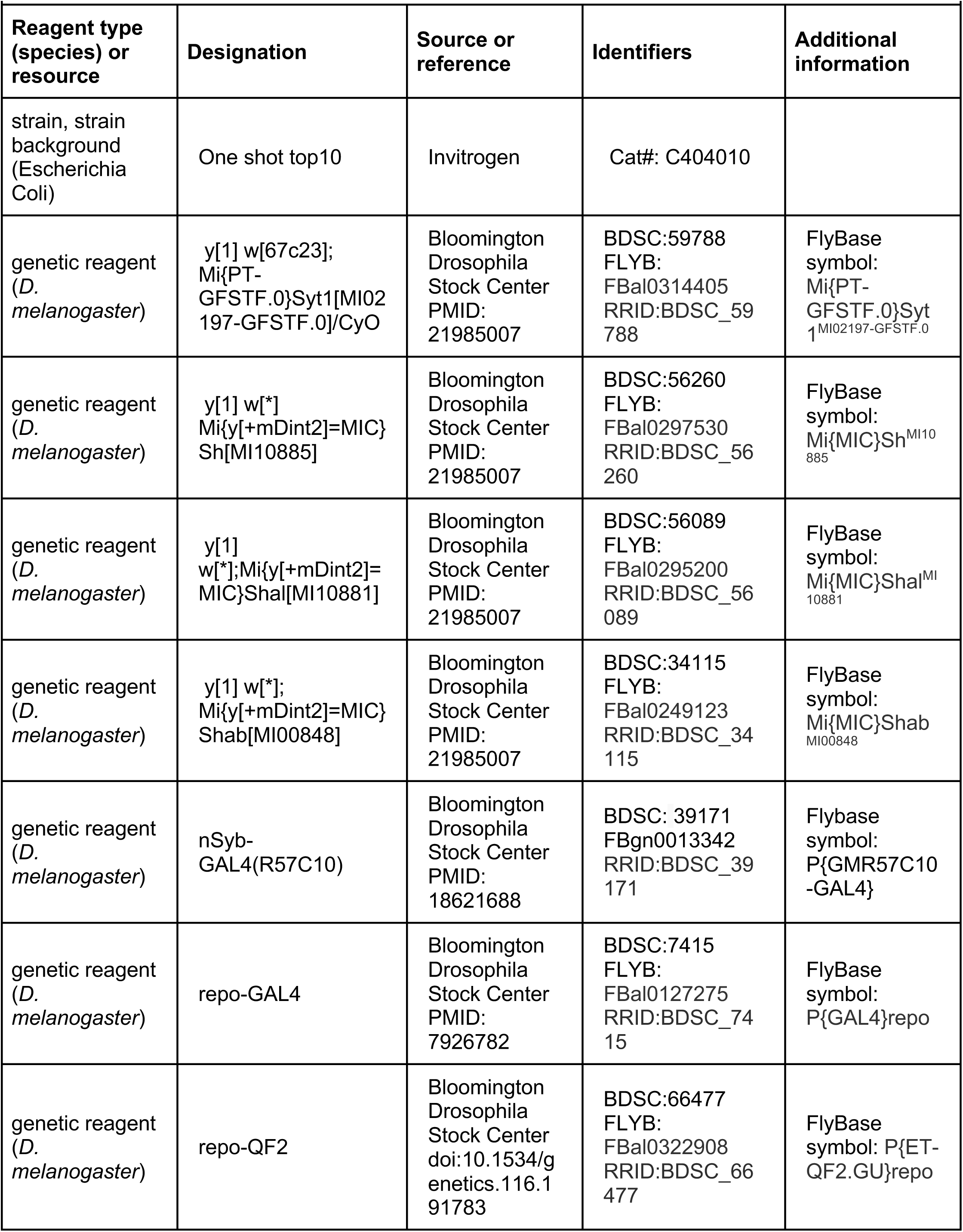
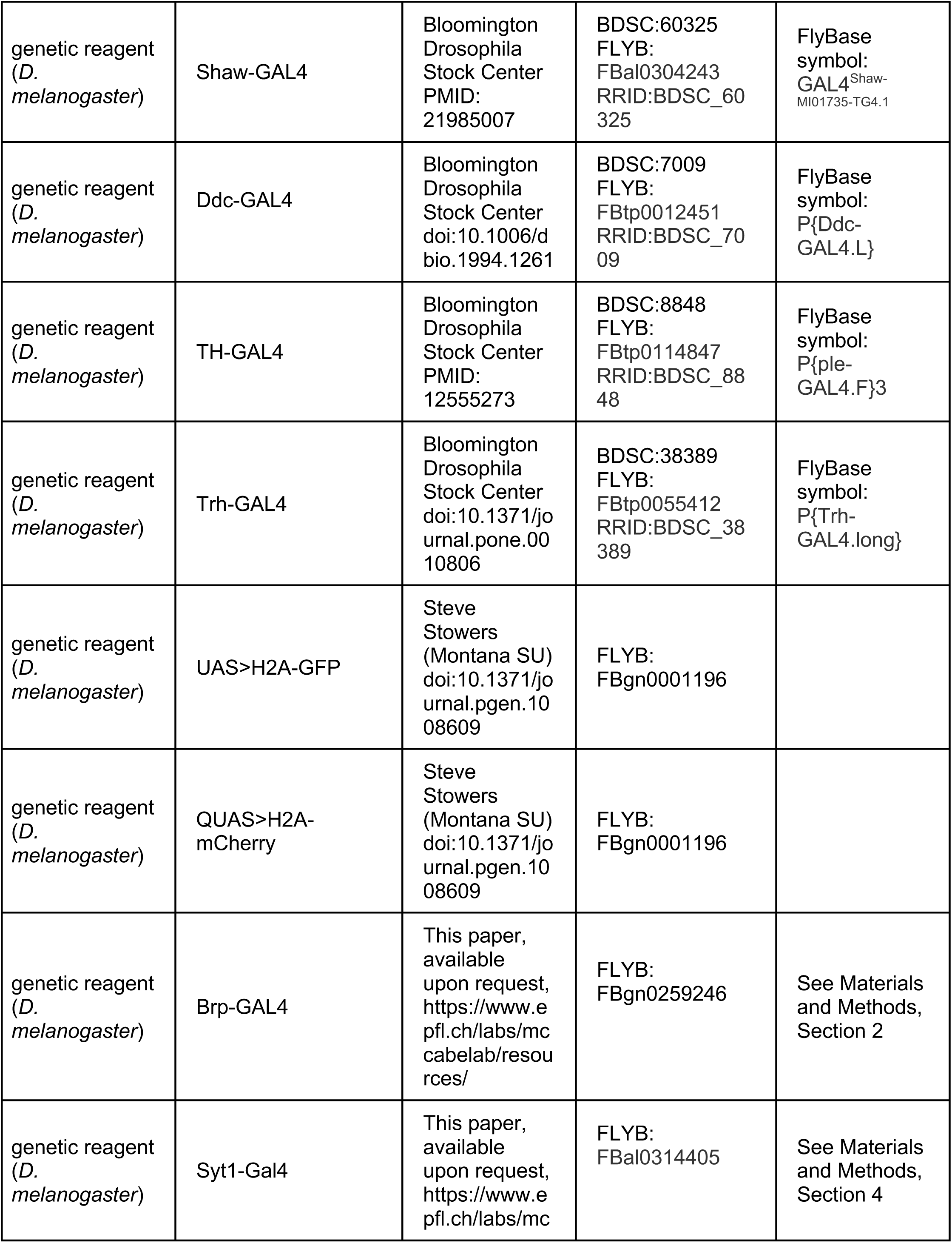
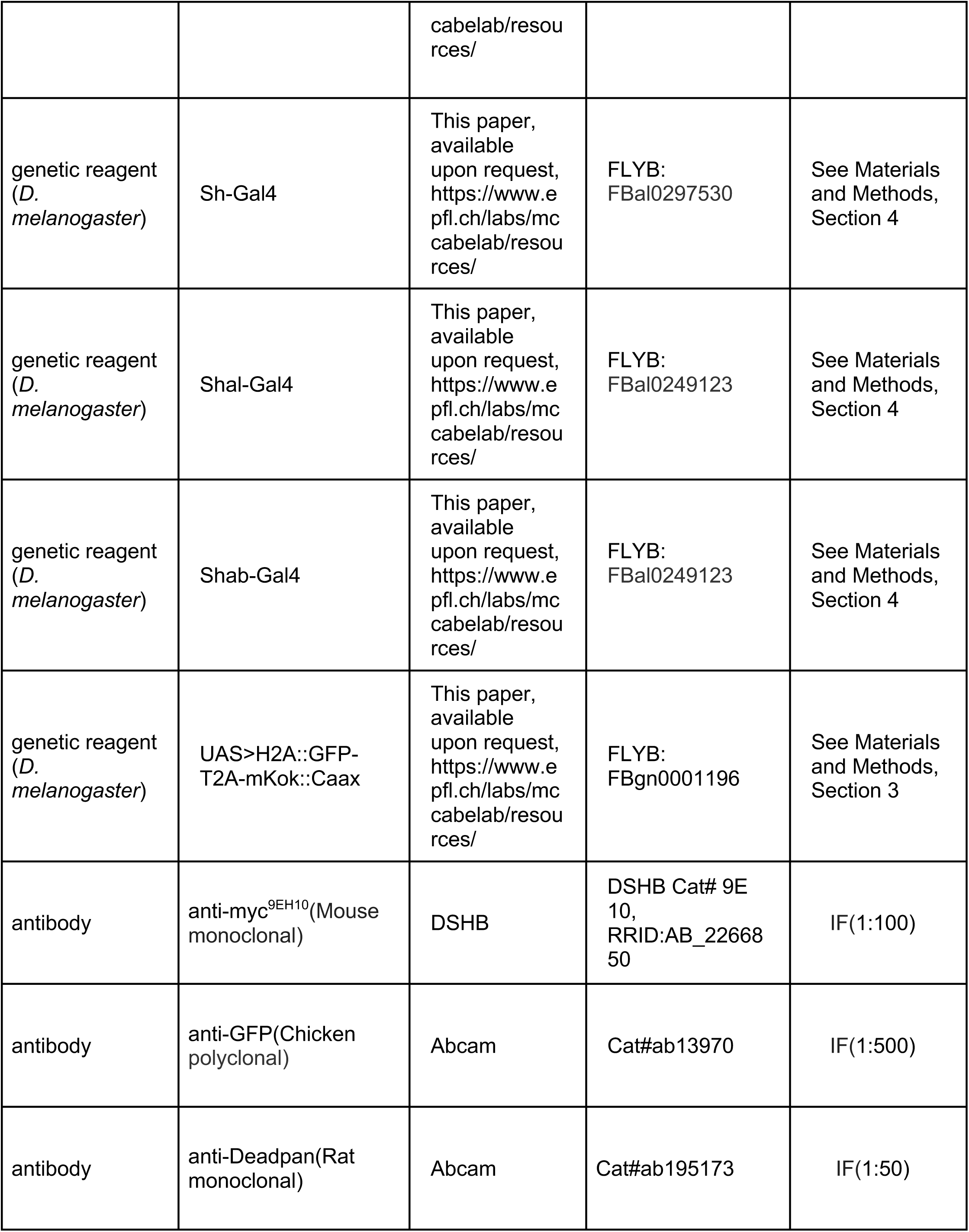
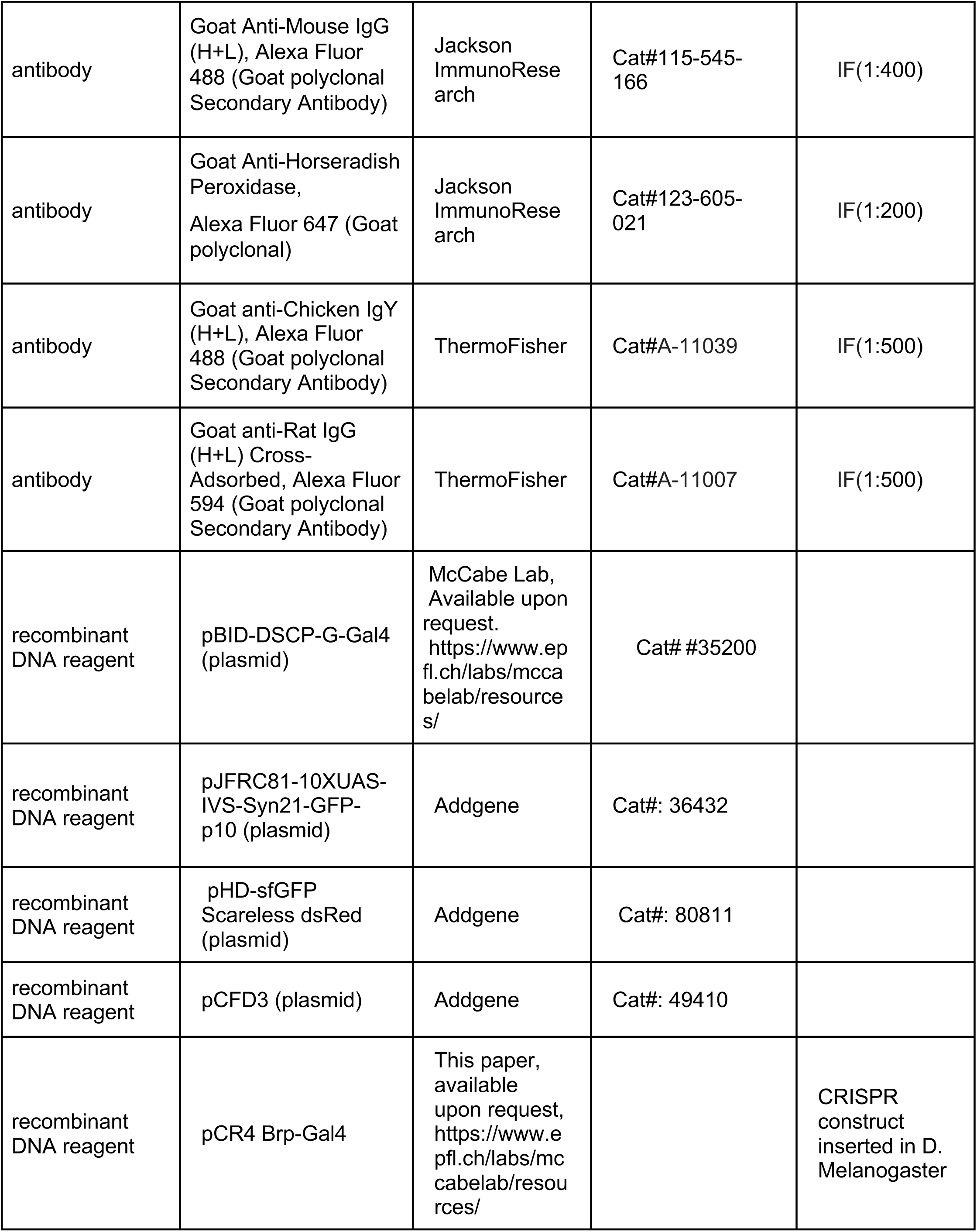
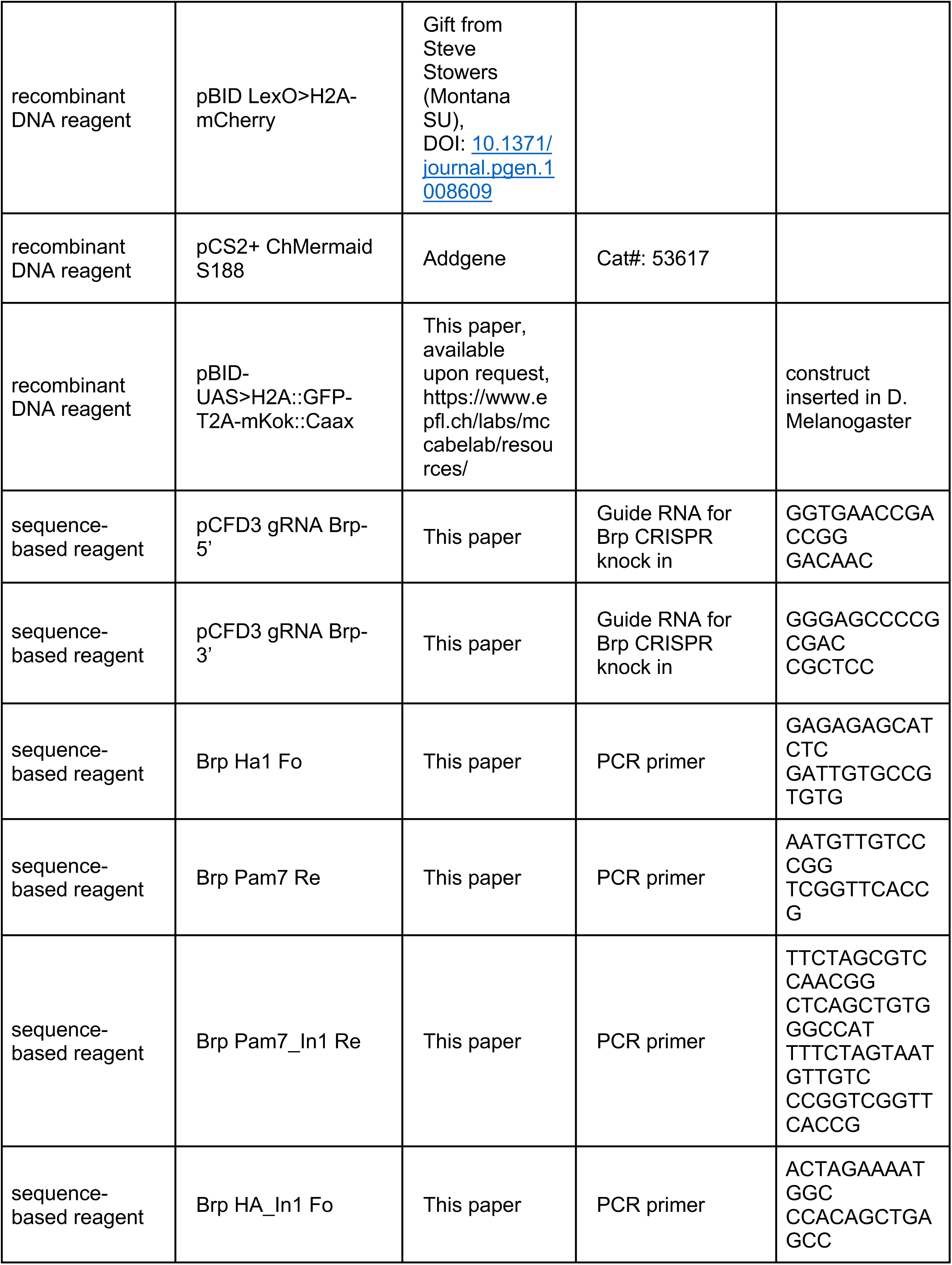
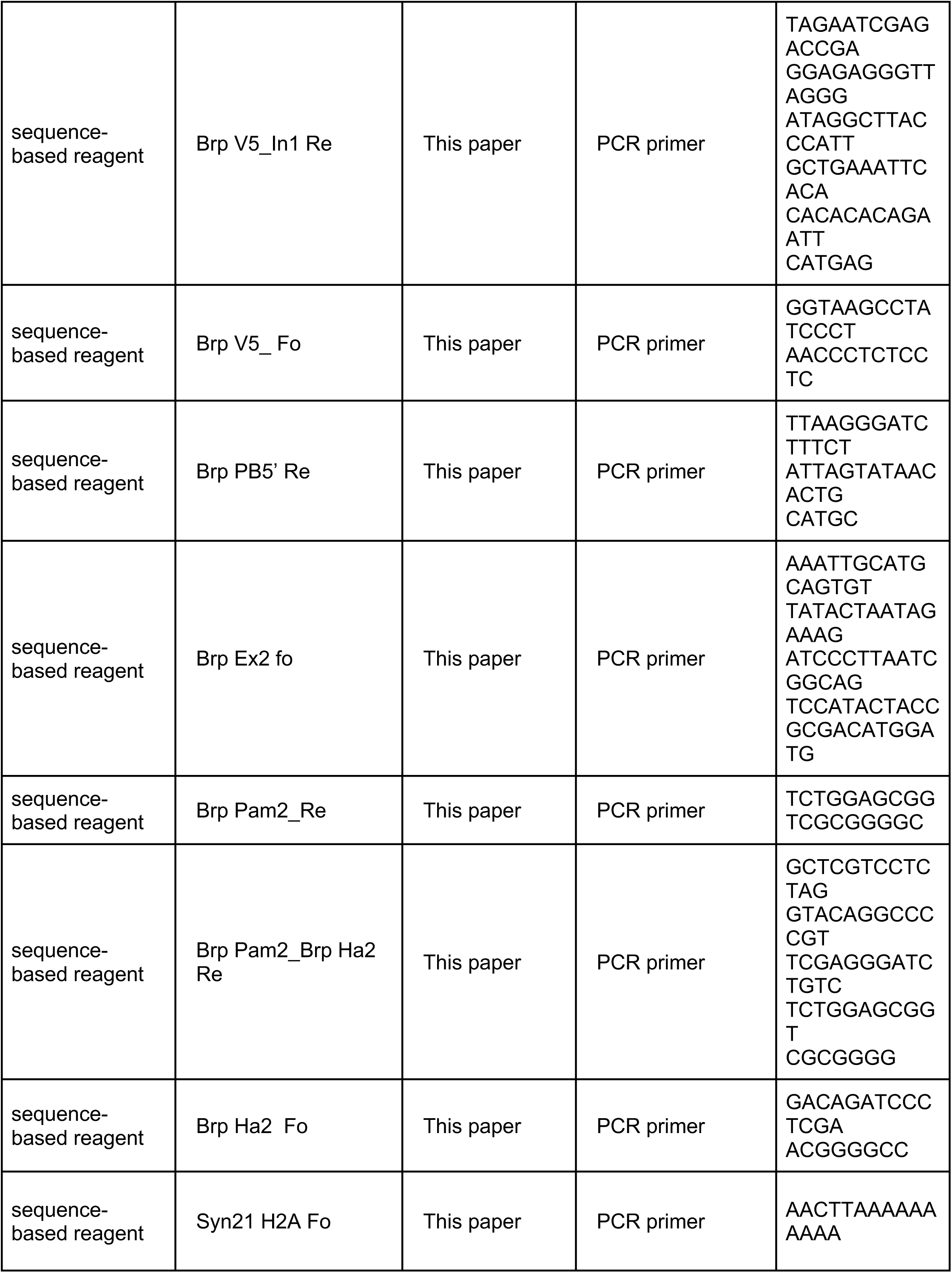
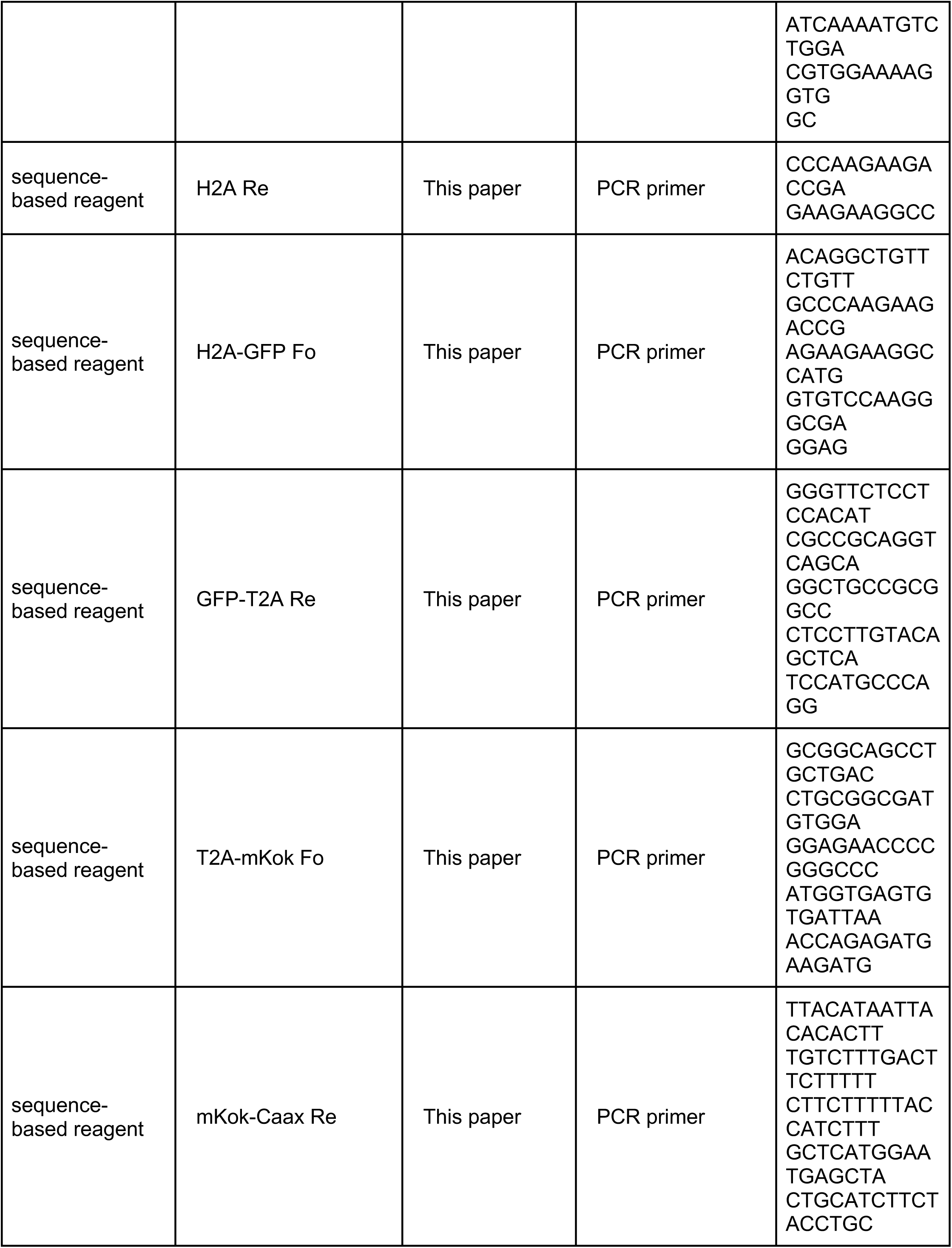
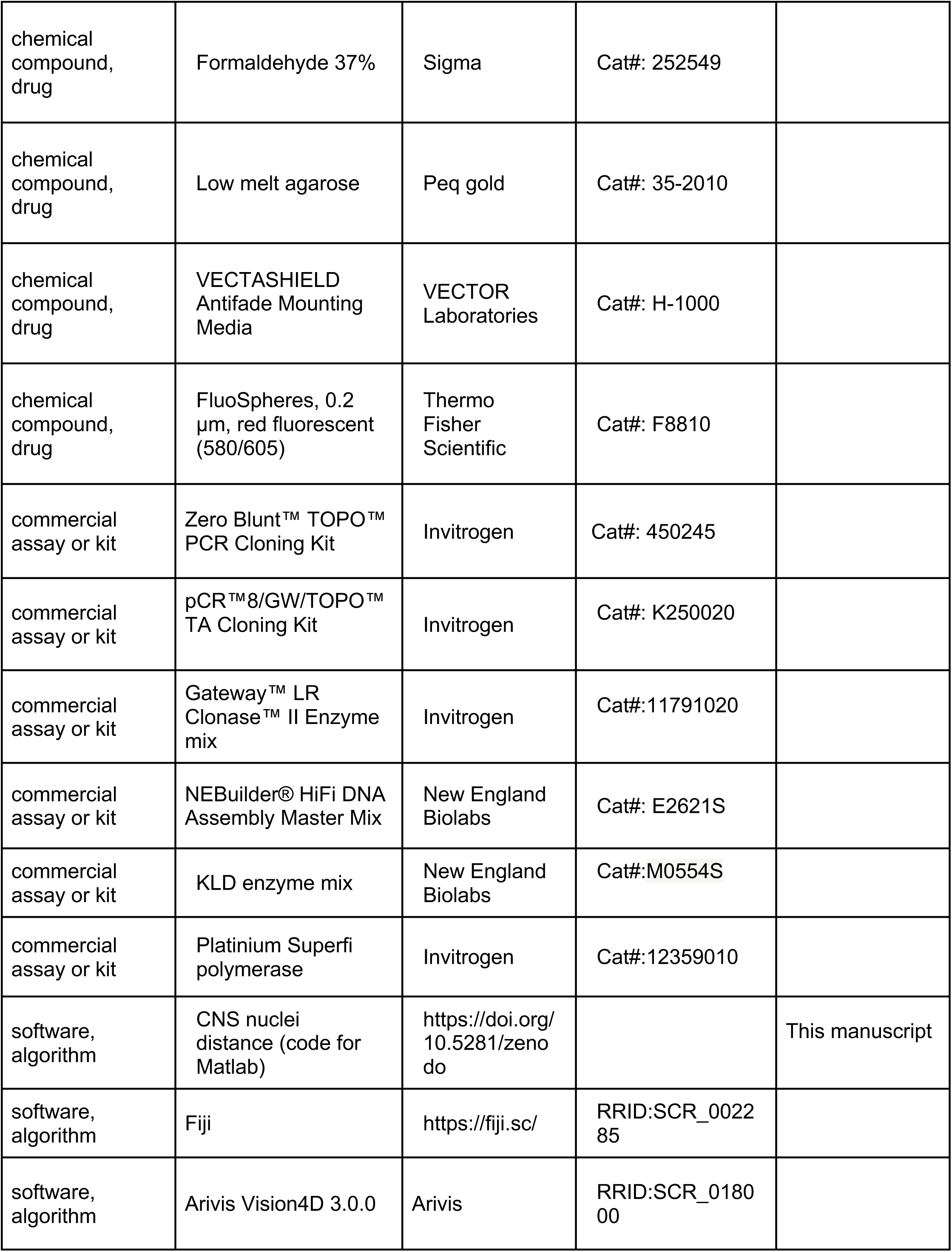
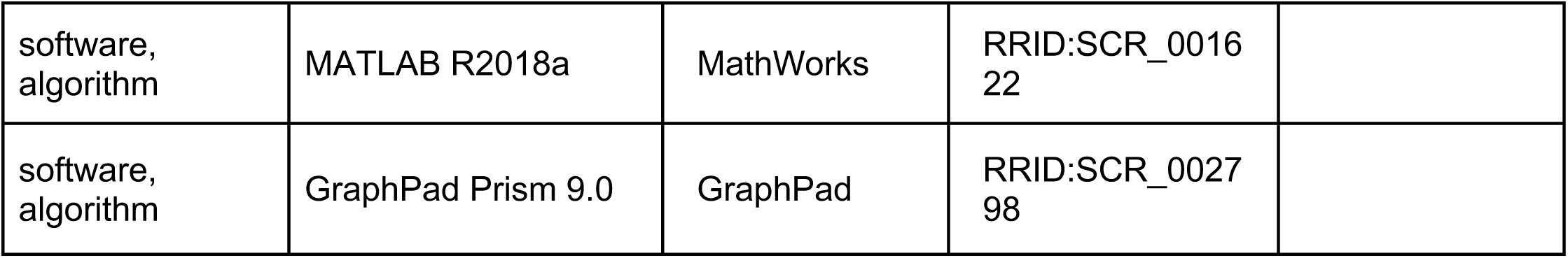

**Figure 2-figure supplement 1.**
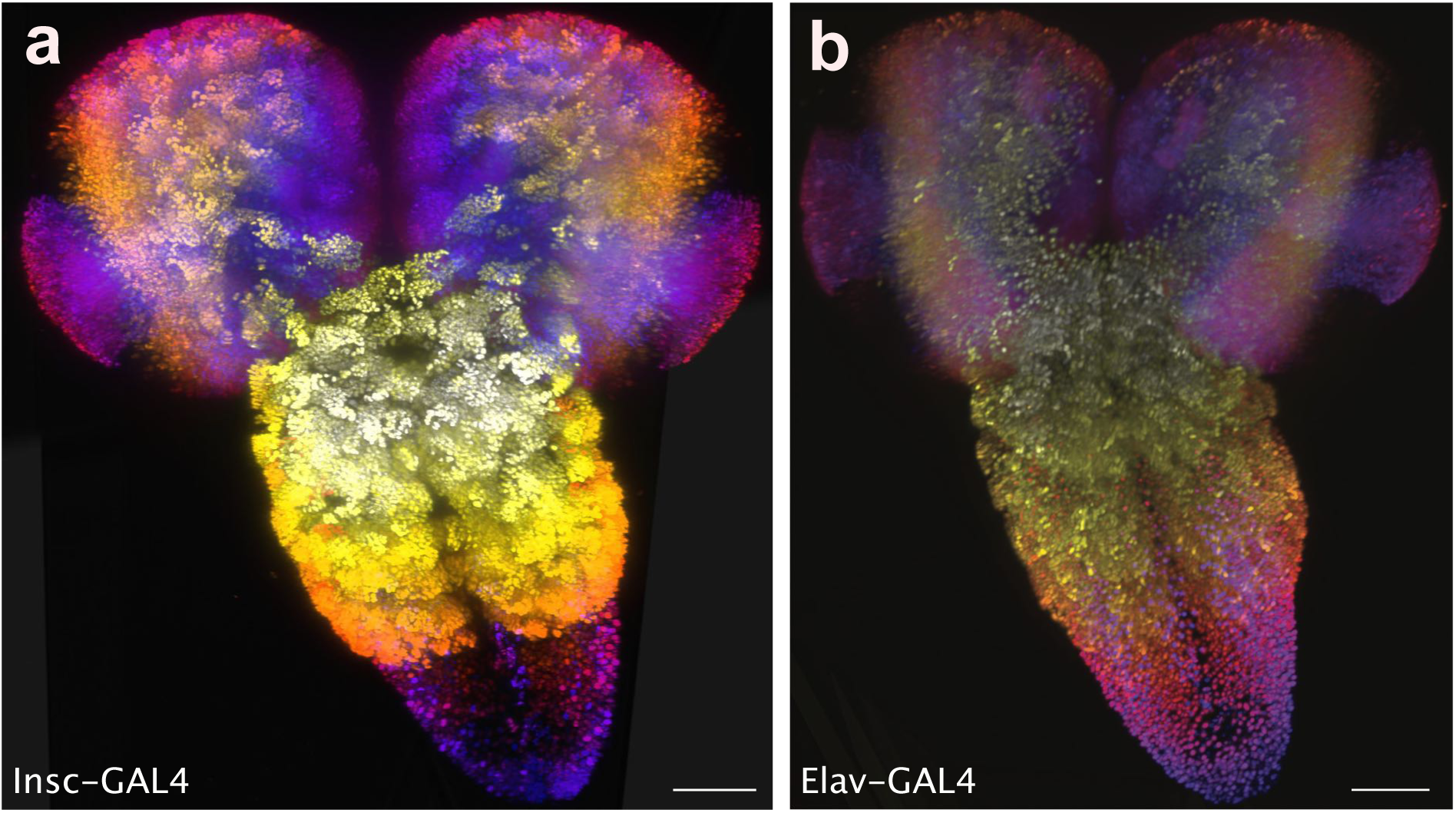
CNS projections labelled with **(a)** Insc-GAL4 and **(b)** Elav-GAL4. Insc-GAL4 labels neuronal stem cells. Elav-GAL4 also labels a fraction of stem cells in addition to potentially glial cells. Colours represent z position. Scale bar: 50μm

**Figure 2-figure supplement 2.**
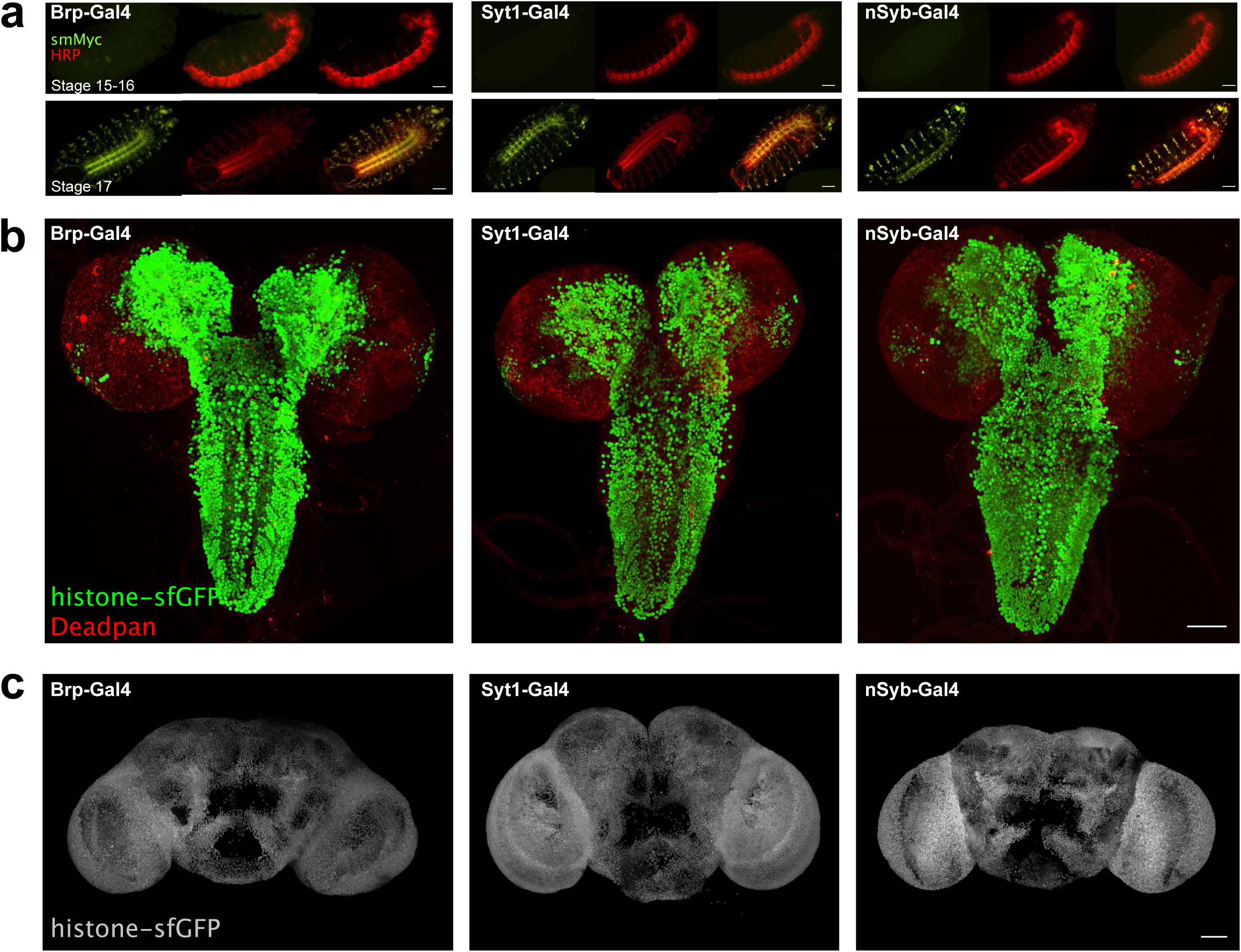
**(a)** Expression UAS>smFP[myc] (green) under control of Brp-Gal4, Syt1-Gal4 and nSyb-Gal4 in embryonic stages 15-16 (upper) and stage 17 (lower). Neuronal membranes are labelled with HRP (red) **(b)** Labeling of L3 larval CNS with Deadpan (red) and UAS>histone-sfGFP (green) under control of Brp-Gal4, Syt1-Gal4 or nSyb-Gal4**(c)** histone-sfGFP expression in the adult brain under control of Brp-Gal4, Syt1-Gal4 and nSyb-Gal4. scale bar = 50μm in **(a)**, **(b)** and **(c)**

**Figure 4-figure supplement 1.**
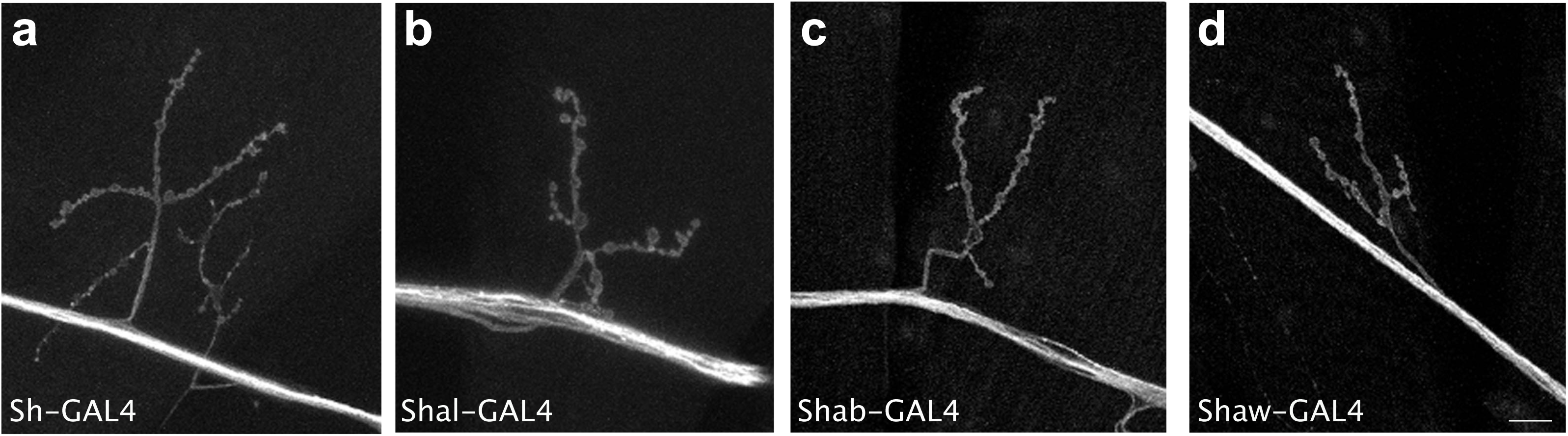
Expression in example motor neurons and neuromuscular junction terminals of UAS>mKO-CAAX by **(a)** Sh-GAL4, **(b)** Shal-GAL4, **(c)** Shab-GAL4 and **(d)** Shaw-GAL4. All four lines express in motor neurons as predicted from mutant analysis. All images identical magnification, scale bar = 10μm.

